# Orally Administered Octenidine Is Effective Against Oral/Throat Pathogens and Safe for the Normal Microbiota of the Oral Cavity, Throat and Large Intestine

**DOI:** 10.1101/2025.07.03.663039

**Authors:** Adam Junka, Malwina Brożyna, Yanfang Sun, Paweł Krzyżek, Michał Tomczyk, Krzysztof Krasucki, Tomasz Matys, Tomasz Musiała, Andrzej Fal

## Abstract

Pharyngitis is a leading cause of outpatient antibiotic use, despite its typically viral or self-limiting nature. Such unnecessary antibiotic therapies are not only the cause of increasing antibiotic resistance, but also significant changes in the human microbiota in the intestines and other locations, which translate into immune disorders and an increased risk of developing several chronic diseases. Orally administered octenidine-containing lozenges offer a topical alternative, but their effects on the host microbiota of the oral cavity, throat, and intestine remain unclear. In this study, we evaluated the antimicrobial and antibiofilm *in vitro* activity of octenidine lozenges against 106 microbial strains, including pathogens and commensals from the oral cavity, pharynx, and large intestine. Minimal biocidal concentrations (MBCs) and minimal biofilm eradication concentrations (MBECs) were determined under physiologically relevant exposure times: 23 minutes for oral contact and 24 hours for intestinal transit. At concentrations achievable in saliva and the intestinal lumen, octenidine effectively eradicated all oropharyngeal pathogens while leaving intestinal commensals unaffected. Its impact on oral commensals resembled that of routine mechanical cleaning. These *in vitro* findings are of high translative value because they support the use of octenidine lozenges as a safe topical treatment for pharyngeal infections - “sore throat”, without adverse effects on the gut microbiota.

## 1. Introduction

The forecasts indicate that by 2050, infections caused by antibiotic-resistant organisms will account for more deaths than cancer and diabetes combined, potentially reaching 10 million fatalities annually (1). While the discovery pipeline for new antibiotics remains narrow, the resistant pathogens continue to proliferate across clinical and community settings. The situation is further exacerbated by the widespread use of antibiotics in agriculture, veterinary medicine, and mass prophylaxis. In many countries, including those with advanced healthcare systems, infections caused by multi-drug-resistant pathogens are already associated with increased morbidity, prolonged hospitalization, higher treatment costs, and limited therapeutic options (2).

While hospital settings receive most attention in discussions on antimicrobial resistance, most of the unnecessary antibiotic consumption and the associated selection pressure occur in ambulatory setting. Antibiotics are frequently prescribed there for self-limiting or non-bacterial infections, such as viral pharyngitis, acute bronchitis, or uncomplicated otitis. This practice, often driven by patient expectations and diagnostic uncertainty, contributes significantly to the emergence and spread of resistant strains in the community. Unlike in hospitals, outpatient antibiotic use is typically less regulated, and treatment durations are often arbitrary, i.e. unnecessarily prolonged or shortened, mostly due to patients’ nonadherence. As a result, large populations may be treated with too low doses of antibiotics, what is one of the main reasons for the development of antibiotic resistance (3). Another factor that complicates antibiotic treatment of infections is the ability of microorganisms to form biofilms—structured communities embedded in the extracellular matrix. Within a biofilm, microbial cells exhibit markedly increased tolerance to antimicrobial agents, elements of the host immune response, and other stressors (4). In the oral cavity and throat biofilm formation on mucosal surfaces, tonsillar crypts, and dental or pharyngeal interfaces is highly relevant and contributes to persistent colonization as well as recurrent infections (5). The oropharynx is a complex microbiological and immunological environment. Its mucosal surfaces are colonized by an extremely diverse community of commensal microorganisms (referred to as the “healthy”, “regular” or “normal” microbiota), which contribute to local defense against hostile strains through niche competition, metabolic exclusion, and immunomodulation (6). These resident species often penetrate deeper into mucosal niches than transient pathogens. Together with the mucosa-associated lymphoid tissue (MALT), the healthy microbiota forms a cooperative defense system that, by outcompeting invaders and by shaping appropriate immune responses, limits pathogen adherence and presence. Disruption of healthy microbiota/pathogens balance - whether by spreading infection or exposure to drugs - can compromise mucosal integrity and immune homeostasis (6).

As mentioned, despite the self-limiting nature of most oropharyngeal infections, antibiotics remain a common first-line intervention. Pharyngitis, tonsillitis, and sore throat are frequently treated empirically with antibiotics (also broad-spectrum ones), even though most cases are viral, non-bacterial in origin. This mismatch between treatment and etiology not only undermines antimicrobial stewardship but also perpetuates the misconception that antibiotics are appropriate for all forms of throat discomfort (7).

Considering these challenges, antiseptic agents (antiseptics) offer a rational alternative for local management of oropharyngeal infections. Unlike antibiotics, they are applied topically, reach high local concentrations, and act through non-specific mechanisms that are difficult for microorganisms to resist, so they do not generate secondary resistance of pathogens to their action (8). Their broad-spectrum activity encompasses bacteria, fungi, and some viruses, making them well-suited for empirical use when the etiological agent is unknown. In the context of sore throat and pharyngitis, antiseptic lozenges alleviate symptoms and reduce microbial load without contributing to the overuse of systemic antibiotics. However, the increasing popularity of antiseptics also raises concerns about their possible overuse and, although unlikely but not excluded, the development of resistance. Documented cases of reduced susceptibility to antiseptic agents such as chlorhexidine - often involving efflux pumps or membrane modifications - highlight that antiseptics are not fully immune to the same evolutionary pressures that compromised antibiotics (9). In contrast to targeted antibiotics, most antiseptics act on multiple microbial structures simultaneously, making the development of high-level, clinically relevant resistance more difficult and less frequent (10). Nevertheless, the growing use of antiseptics across medical, cosmetic, and consumer products requires careful monitoring. Their value as part of antimicrobial stewardship depends not only on efficacy, but also on judicious and evidence-based application. Thus, integrating antiseptics into routine management of oropharyngeal complaints offers a pragmatic approach to slowing development of antibiotic resistance.

One of such antiseptics that can be applied in that context is referred to as the octenidine dihydrochloride, abbreviated as “OCT”, i.e. N,N′-(1,10-decanediyldi-1(4H)-pyridinyl-4-ylidene)bis(1-octanamine) dihydrochloride) bispyridine cationic antiseptic that has been already used for skin, mucosal, and wound antisepsis (**Figure 1**).

**Figure 1.**
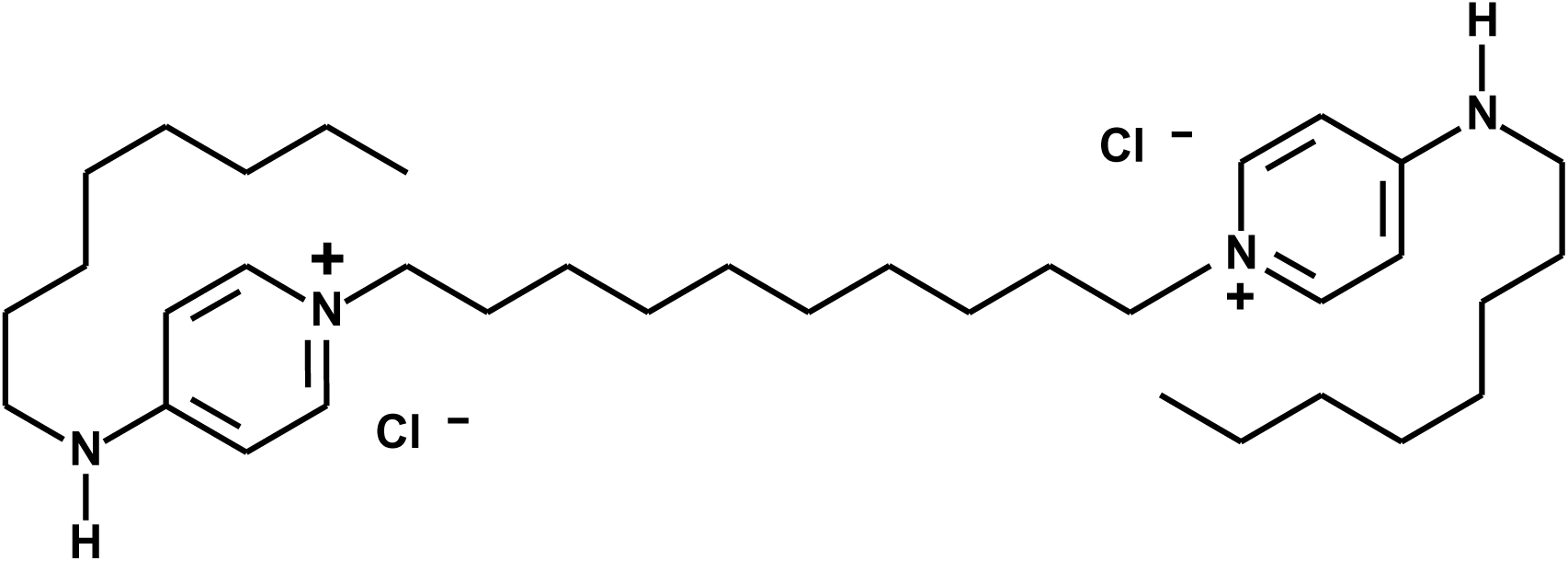
Chemical structure of octenidine dihydrochloride (N,N′-(1,10-decanediyldi-1(4H)- pyridinyl-4-ylidene)bis(1-octanamine) dihydrochloride.

It exhibits broad-spectrum antimicrobial activity against Gram-positive and Gram-negative bacteria, fungi, and some enveloped viruses. Its mechanism of action involves disruption of microbial membranes, leading to rapid cell death without relying on specific intracellular targets (11,12). Octenidine has a favorable safety profile, low/no systemic absorption, and minimal tissue irritation, making it suitable for applications involving mucosal surfaces (13). Although its use has been well established in external topical formulations, the potential of octenidine for oral administration - particularly in lozenge form- has only recently begun to attract clinical and scientific interest. In our recent study (14), we demonstrated that octenidine lozenges exhibit potent biocidal activity against a broad range of oropharyngeal pathogens, including *Streptococcus pyogenes*, *Staphylococcus aureus* and *Candida albicans*. The formulation was shown to dissolve completely within an average of 23 minutes, ensuring prolonged contact with the mucosal surface and sustained local exposure to the antiseptic agent. Pharmacokinetic evidence confirms that octenidine dihydrochloride is virtually not absorbed through gastrointestinal mucosa or the skin, passing intact through the gastrointestinal tract and exiting in feces (13). Both the European Public Maximum Residue Limits (MRL) Assessment Report and DrugBank note its negligible systemic uptake - “virtually not absorbed”—making significant systemic or intraluminal concentrations highly unlikely (15,16).

Antiseptic lozenges offer several practical and pharmacodynamic advantages. They are easy to administer, portable, and quite acceptable to patients outside clinical settings. Most importantly, they provide sustained release and prolonged mucosal exposure, which enhances antimicrobial efficacy without requiring patient supervision (17). This gives a completely different quality than mouthrinses that contact the oropharyngeal mucosa for less than a minute, lozenges can maintain antiseptic presence in saliva and on the mucosa for over a quarter of an hour. This extended contact time is particularly valuable in targeting pathogens embedded in biofilms or located in deeper epithelial structures.

Despite these advantages, the oral administration of antiseptics raises two important safety concerns that have not been adequately addressed yet. The first relates to the potential impact of swallowed antiseptic on the gut microbiota, particularly in the large intestine, where microbial diversity plays a key role in health and immune regulation (18). Given that disturbances in intestinal microbiota can contribute to conditions such as dysbiosis or antibiotic-associated diarrheas related to the overgrowth of *Clostridioides difficile*, assessing the gut safety of orally administered antiseptics is clinically relevant (19).

The second concern involves the local effect in the oropharynx: whether antiseptics indiscriminately eliminate not only pathogens but also commensal microorganisms that are essential for maintaining mucosal homeostasis. Commensal microorganisms such as *Streptococcus salivarius*, *Streptococcus mitis*, or *Limosilactobacillus spp.* play important roles in colonization resistance, immune modulation, and maintenance of mucosal health (20). If antiseptics eliminated beneficial species along with pathogens, they could inadvertently disrupt the local microbial balance and weaken host defense mechanisms. Despite the growing interest in orally administered octenidine-containing lozenges, no studies have compared the susceptibility of commensal and pathogenic strains in both planktonic and biofilm forms to conditions mimicking physiological contact times in the mouth and gut. Lack of these data leaves clinicians without evidence needed to evaluate the microbiological safety of antiseptics and their potential impact on host-associated microbial communities. Therefore, in this study, we evaluated the antimicrobial and antibiofilm activity of octenidine dihydrochloride against a panel of clinical and reference strains representing both pathogenic and commensal microorganisms of the oral cavity, pharynx, and colon. This study aims to provide translationally relevant *in vitro* data on the use of octenidine-containing lozenges, with particular focus on their antimicrobial efficacy against pathogens and their safety represented by little or no effect on the physiological microbiota.

## 2. Material and Methods

### 2.1. Antiseptic agent, strains and culturing conditions

The octenidine dihydrochloride used in this study was obtained either as analytical-grade powder (≥98% purity; Thermo Fisher Scientific, Waltham, MA, USA) or in the form of commercially available lozenges containing 2.6 mg of octenidine dihydrochloride per unit (total lozenge mass = 832 mg), provided in sealed blister packs under the brand name Octeangin® (Klosterfrau, Cologne, Germany, LOT:178123). Following initial comparative tests, lozenge-derived octenidine was selected for MBC (minimal biocidal concentration) and MBEC (minimal biofilm eradication concentration) analyses, as this formulation better reflects the clinically relevant route of administration investigated in the present study. A total of 106 clinical and reference strains were used in this study, from the internal collection of the P.U.M.A. (Platform for Unique Models Application) and Zhejiang International Joint Laboratory of Traditional Medicine and Big Health Products Development. The tested microorganisms included: *Staphylococcus aureus* MRSA (methicyllin-resistant *Staphylococcus aureus*) (n = 15), *Streptococcus pyogenes* (n = 15), *Escherichia coli* (non-pathogenic isolates, n = 15), *Candida albicans* (n = 15), *Lactobacillus reuteri* (n = 15), *Moraxella catharralis* (n = 10), *Streptococcus salivarius* (n = 5), *Streptococcus mitis* (n = 5), *Haemophilus spp.* (n = 5 species: *parainfluenzae* = 3 and *haemolyticus* = 2), and *Bacteroides* spp. (n = 5, species: *thetaiotaomicron* = 3; *fragilis* = 2). The reference (ATCC-American Type Culture Collection, Manassas, VA, USA) strains applied were *S. aureus* 29213, *Str. Pyogenes* 12344, E. *coli* 25922, *C. albicans* 1021, *L. reuterii* 23272, *M. catharalis* 25238 and only for modified disk diffusion method, the *Pseudomonas aeruginosa* 27853 was applied. All clinical strains were isolated from patient samples in accordance with institutional guidelines and with approval from the Wroclaw Bioethical Committee (KB949-2022). Frozen glycerol stocks (−80 °C) were revived by streaking onto appropriate solid media under optimal atmospheric conditions. Aerobic bacteria (e.g., *S. aureus, E. coli*, *S. pyogenes*) were cultured on tryptic soy agar (TSA; Biomerieux, <u>Marcy-l’Étoile</u>, France) or Columbia agar with 5% sheep blood (Biomerieux, <u>Marcy-l’Étoile</u>, France) at 37 °C. Anaerobic species (*Bacteroides spp*.) were cultured on Schaedler agar supplemented with hemin and vitamin K1 (Biomerieux, <u>Marcy-l’Étoile</u>, France) and incubated at 37°C in anaerobic workstation under an atmosphere of 85% N₂, 10% H₂, and 5% CO₂. Microaerophilic organisms (*Haemophilus spp., Moraxella spp.)* were cultured on chocolate agar (Biomerieux, <u>Marcy-l’Étoile</u>, France) or Haemophilus-selective media (Thermo Fisher Scientific, Waltham, MA, USA) under microaerophilic conditions at 37°C (CampyGen, Oxoid, UK). *Candida albicans* were maintained on Sabouraud dextrose agar (SDA; Biomerieux, <u>Marcy-l’Étoile</u>, France) at 35 °C under aerobic conditions. After 18–48 hours of incubation (species-dependent), colonies were evaluated for purity based on morphological features. Representative colonies were further examined microscopically by Gram staining and lactophenol blue for *C. albicans*) to confirm expected morphology and eliminate potential contaminants. Only morphologically appropriate and Gram-consistent colonies were selected for experimental use. All strains were subcultured no more than twice prior to susceptibility testing to preserve phenotypic stability.

### 2.2. Estimation of Octenidine Concentration Along the Gastrointestinal Tract

Following complete dissolution of one lozenge containing 2.6 mg of octenidine dihydrochloride in the stomach (assumed volume of 200 mL) (14), the resulting gastric concentration was calculated to be approximately 13 mg/L. Upon gastric emptying, this content enters the small intestine and becomes progressively diluted by endogenous secretions. On average, approximately 2 L of digestive fluids—including bile, pancreatic juice, and gastric secretions—are secreted into the small intestine daily, resulting in a further dilution of the antiseptic to approximately 1.18 mg/L in the small intestinal lumen (21). As the intestinal content transitions to the large intestine, it mixes with approximately 1 L of colonic fluid, representing the aqueous fraction of fecal matter (22). This final step leads to a calculated colonic concentration of ∼0.05 mg/L. This value was therefore selected as the reference exposure level for in vitro testing of octenidine’s impact on representative members of the intestinal microbiota. Our model was deliberately constructed to overestimate potential colonic concentrations of biologically active octenidine. Several factors known to reduce antiseptic availability in vivo—including chemical and enzymatic degradation, adsorption to mucins, dietary components, and microbial biomass—were not accounted for in the present calculation. Consequently, the adopted colonic concentration of 0.05 mg/L should be interpreted as a conservative upper-limit estimate.

### 2.3. Initial indication of antimicrobial activity of octenidine-containing lozenges using a modified disk diffusion method

For this purpose, three pharmacopoeial reference strains were used as representatives of Gram-negative bacteria, Gram-positive bacteria, and fungi: *P. aeruginosa* ATCC 27853, *S. aureus* ATCC 29213, and *C. albicans* ATCC 10231, respectively. Overnight cultures were adjusted to a 0.5 McFarland standard (∼2 × 10⁸ CFU/mL (colony-forming unit) for bacteria and ∼2 × 10⁶ CFU/mL for *C. albicans*) using a densitometer (Den-1, Biosan, Piła, Poland). Standardized microbial suspensions were then spread evenly onto the surface of Mueller-Hinton Agar (MHA; Biomaxima, Lublin, Poland) poured into 90 mm diameter Petri dishes (Noex, Komorniki, Poland). An 8 mm diameter well was created at the center of each agar plate using a cork borer (Equimed, Kraków, Poland), into which a single octenidine-containing lozenge was placed. Plates were incubated for 24 hours at 37 °C in a microbiological incubator (Binder, VWR International LLC, Radnor, PA, USA). After incubation, the area of microbial growth inhibition and partially dissolved lozenge was measured (in mm²) using ImageJ software (National Institutes of Health, Bethesda, MD, USA; https://imagej.nih.gov/ij/). The factual area of microbial growth inhibition (mm^2^) was calculated by subtracting the later mentioned parameter from the earlier mentioned parameter. Control plates containing inoculated MHA without lozenges served as growth controls. All experiments were performed in triplicate.

### 2.4. Determination of tested strains’ ability to form biofilm in vitro using crystal violet method

The biofilm-forming capacity of each strain was evaluated using a standardized crystal violet (CV) assay in flat-bottom 96-well polystyrene (Wuxi Nest Biotechnology, Wuxi, Jiangsu, China) microtiter plates. For bacterial strains, overnight cultures were adjusted to 0.5 McFarland standard (∼2 × 10⁸ CFU/mL), while for *Candida albicans*, a 0.5 McFarland suspension corresponded to ∼2 × 10⁶ CFU/mL, using a densitometer (Den-1, Biosan, Piła, Poland). Each suspension was diluted 1:100 in the appropriate growth medium to yield a final inoculum of approximately 1 × 10⁶ CFU/mL for bacteria and ∼1 × 10⁴ CFU/mL for *C. albicans*. Aliquots of 200 µL were transferred into individual wells (n = 3 per strain). Plates were incubated under strain-specific atmospheric conditions to allow for biofilm formation: aerobic bacterial strains (e.g., *S. aureus*, *E. coli*, *S. pyogenes*) and *C. albicans* were incubated at 37 °C under ambient air for 24 h; microaerophilic species (e.g., *Haemophilus* spp., *Moraxella catarrhalis*) were incubated for 48 h using CampyGen™ gas packs (Oxoid, UK); anaerobic strains (*Bacteroides spp*.) were incubated for 72–96 h at 37°C in an anaerobic workstation (Whitley A35, Don Whitley Scientific, UK) under a controlled atmosphere of 85% N₂, 10% H₂, and 5% CO₂. Appropriate growth media were used for each species, as previously described for strain preparation, including Schaedler agar/broth (anaerobes), chocolate or *Haemophilus*-selective media (microaerophiles), and Sabouraud dextrose medium for *C. albicans*.

Following incubation, non-adherent (planktonic) cells were gently aspirated from each well, and the wells were rinsed twice with 200 µL of sterile phosphate-buffered saline (PBS; Thermo Fisher Scientific, Waltham, MA, USA) to remove residual non-adherent biomass. The remaining surface-attached biofilm was stained with 0.1% crystal violet solution (Chempur, Piekary Slaskie, Poland) (200 µL per well) for 15 minutes at room temperature. Excess stain was removed by washing with distilled water 3x/200 µL, and plates were air-dried. Bound crystal violet was then solubilized with 200 µL of 96% ethanol (Chempur, Piekary Slaskie, Poland), and absorbance was measured at 590 nm using a microplate reader (Multiskan™ FC, Thermo Fisher Scientific, Waltham, MA, USA). All experiments were performed in technical triplicate and biological duplicate and indicated as the average value.

### 2.5. Determination of Minimal Biocidal Concentration using microdilution method in 96-well plates

MBCs of octenidine dihydrochloride (analytical-grade powder) were determined using a modified broth microdilution protocol adapted from Clinical and Laboratory Standards Institute (CLSI) guidelines (M26-A, M27-A4 for yeasts), and adjusted to account for the physicochemical properties of the antiseptic agent. For each bacterial strain, a suspension equivalent to 0.5 McFarland standard (approximately 2 × 10⁸ CFU/mL) was prepared (using a densitometer) in the appropriate growth medium and diluted 1:100 to yield a final inoculum of ∼1 × 10⁶ CFU/mL. For *Candida albicans*, a 0.5 McFarland suspension corresponded to ∼2 × 10^4^ CFU/mL. A 100 µL aliquot of these suspensions were added to each well of a sterile 96-well microtiter plate containing 100 µL of octenidine dihydrochloride, serially diluted in two-fold geometric progression across a range of final concentrations from 250 mg/L to 0.05 mg/L. Plates were incubated under species-appropriate conditions: aerobic strains were maintained at 37 °C in ambient air; anaerobic strains were incubated in an anaerobic chamber at 37 °C; and microaerophilic organisms were cultured using CampyGen gas packs also at 37°C The incubation time was set to 23 minutes for strains typically colonizing the oral cavity and throat, reflecting the average contact time of orally administered octenidine. For strains representing the intestinal microbiota, the incubation time was extended to 24 hours to simulate the estimated retention and exposure time of octenidine in the large intestine. Following incubation, the entire contents (200 µL) of each well showing no visible turbidity were transferred into 1.8 mL of validated neutralizing solution composed of phosphate-buffered saline (PBS) supplemented with 3% Tween 80, 0.3% lecithin, 0.1% histidine, 0.1% cysteine, and 0.5% sodium thiosulfate. After 10 minutes of incubation at room temperature for antiseptic inactivation, 100 µL of each neutralized sample was inoculated into 5 mL of fresh, sterile broth corresponding to the organism’s original growth medium and incubated again under the same atmospheric conditions. Absence of visible growth after 24–48 hours was interpreted as evidence of biocidal activity at the tested concentration and the MBC value was determined at the minimal of these concentrations. All experiments were performed in duplicate, and the modal MBC value was recorded for each strain.

### 2.6. Determination of Minimal Biofilm Eradication Concentration (MBEC)

MBECs of octenidine dihydrochloride (analytical-grade powder) were determined using a modified static biofilm protocol based on the broth microdilution method in 96-well flat-bottom microtiter plates. For each strain, an inoculum of approximately 1 × 10^4^ CFU/mL was prepared as described above and seeded into sterile plates (200 µL per well). Plates were incubated under species-appropriate atmospheric conditions (as described in the section Antiseptic agent, strains and culturing conditions) to allow biofilm formation: 24 h for aerobic organisms, 48 h for microaerophilic species, and 72–96 h for anaerobic strains, depending on their individual growth kinetics. Following the biofilm growth phase, planktonic cells were gently aspirated from each well, and biofilms were rinsed once with 200 µL of sterile phosphate-buffered saline (PBS). Fresh medium containing octenidine dihydrochloride in serial two-fold dilutions (250– 0.05 mg/L) was then added (200 µL per well), and plates were re-incubated under the same atmospheric conditions for 23 minutes or 24 hours to simulate the estimated retention and exposure time of octenidine in the oral cavity/throat or large intestine, respectively. After antiseptic exposure, the entire contents of each well were aspirated and transferred into 1.8 mL of validated neutralizing solution (PBS with 3% Tween 80, 0.3% lecithin, 0.1% histidine, 0.1% cysteine, and 0.5% sodium thiosulfate). During aspiration, the biofilm was deliberately resuspended by intensive pipetting (aspirating and dispensing at least three times per well) to ensure maximum recovery of viable cells. Additionally, neutralizing solution (200 µL) was added directly to the residual material remaining in the wells, removed, and after 10 minutes of inactivation, 200 µL of fresh growth medium was added to each well. All resuspended samples (from neutralized suspensions and directly treated well contents) were incubated for up to 72 hours under original atmospheric conditions. The absence of visible growth in both the transferred suspension (Falcon tubes or microcentrifuge tubes) and in the original plate wells was considered as indicative of complete biofilm eradication at the tested concentration and the MBEC value was determined at the lowest of these concentrations. All assays were conducted duplicate, and the modal MBEC value was recorded for each strain.

### 2.7. Statistical Analysis

All numerical data were analyzed using GraphPad Prism (version 10.4.2, GraphPad Software, San Diego, CA, USA). The normality of data distribution was assessed using the Shapiro–Wilk test. For datasets with normal distribution, group means were compared using one-way analysis of variance (ANOVA). The assumption of homogeneity of variances was verified using Brown– Forsythe and Bartlett’s tests. As no significant variance inequality was detected, Tukey’s multiple comparison test was applied post hoc. A p-value < 0.05 was considered statistically significant.

### 2.8. Chemical formula drawing

The formula of octenidine dihydrochloride was drawn using ChemWindow software (Bio-Rad Laboratories, Inc., Hercules, CA, USA).

## 3. Results

The antimicrobial activity of the lozenges towards three references, pharmacopeia-relevant strains of Gram-positive, Gram-negative bacteria and fungi, was confirmed using a modified disk diffusion method, serving as a prerequisite experiment for subsequent analyses (**Figure 2**, **Table 1**) As seen in **Figure 2**, clear zones of growth inhibition were observed around the octenidine-containing lozenges for all tested reference strains (*S. aureus*, *P. aeruginosa*, *C. albicans*), confirming their antimicrobial activity. The extent of inhibition varied between species, with the largest zone recorded for *C. albicans* (growth inhibition covering whole plate), followed by *S. aureus*, and the smallest for P*. aeruginosa* (mean area: 2957±162 mm²), as shown in **Table 1**. Control plates showed confluent microbial growth in the absence of lozenges, confirming the validity of the model. These results supported the use of lozenges in subsequent MBC and MBEC testing.

**Figure 2.**
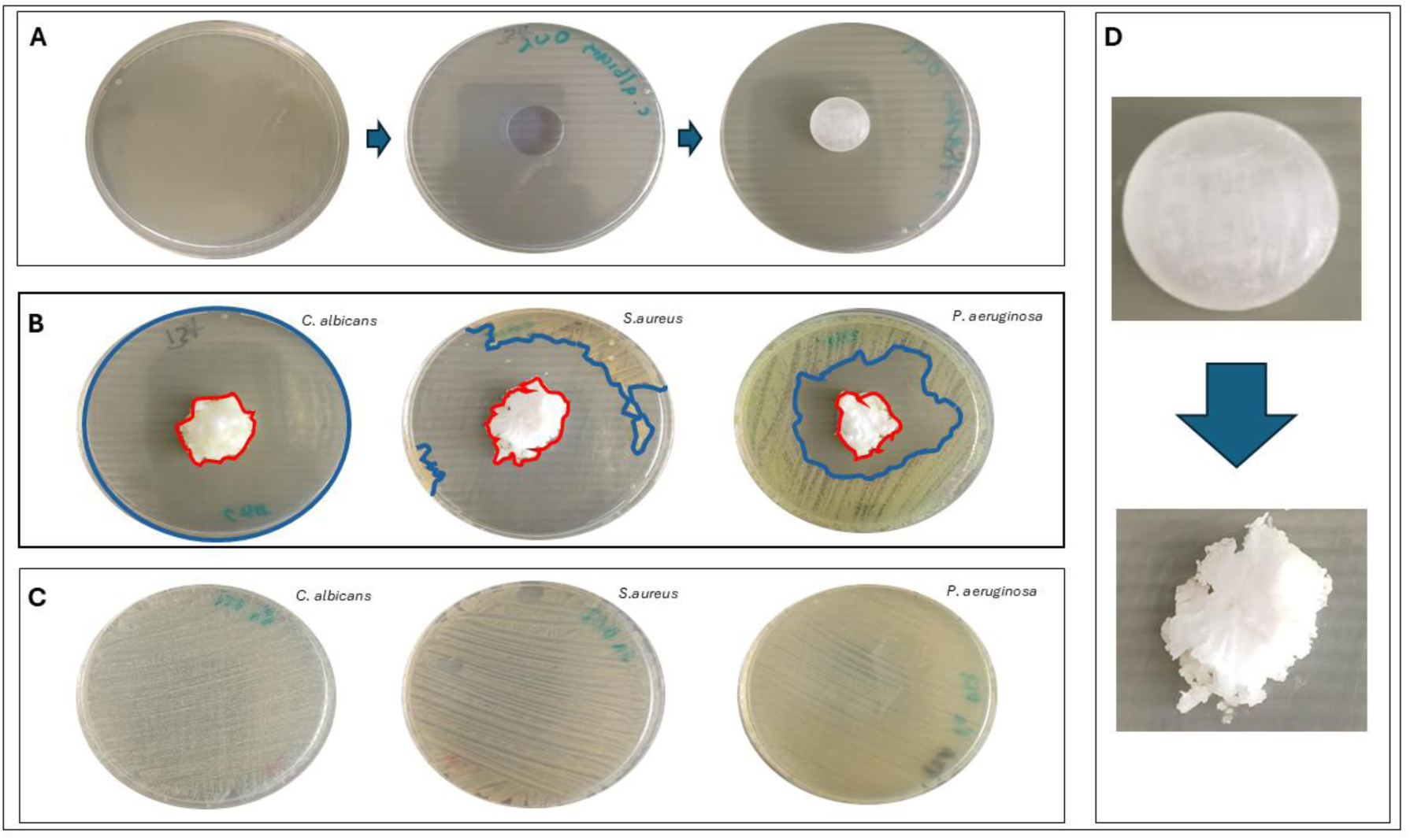
Assessment of the antimicrobial activity of octenidine-containing lozenges using a modified disk diffusion method. **A** – preparation steps of a Mueller-Hinton Agar (MHA) plate inoculated with microorganisms and containing an octenidine lozenge. **B** – measurement of microbial growth inhibition zones (mm²) using ImageJ software; the red line marks the partially dissolved lozenge (mm^2^), while blue lines delineate the inhibition area (mm^2^). **C** – microbial growth control without lozenge. **D** – visualization of the lozenge during partial dissolution on the agar surface. Plate diameter is 90mm and the surface area is equal to 6361.7 mm^2^.

**Table 1.** Area (mm^2^) of microbial growth inhibition zone of pharmacopeial reference strains.

| Area of microbial growth inhibition (mm<sup>2</sup>) |  |  |
| --- | --- | --- |
| *C. albicans* | *S. aureus* | *P. aeruginosa* |
| 5540±51 | 5004±74 | 2957±162 |

Next, the biofilm-forming ability of the tested strains (n=105) was assessed using crystal violet assessment method (**Figure 3**). While all strains demonstrated the capacity to form biofilm *in vitro*, significant, inter- and intra-species-related differences were observed in the quantity of biofilm biomass produced (**Figure 4**.)

**Figure 3.**
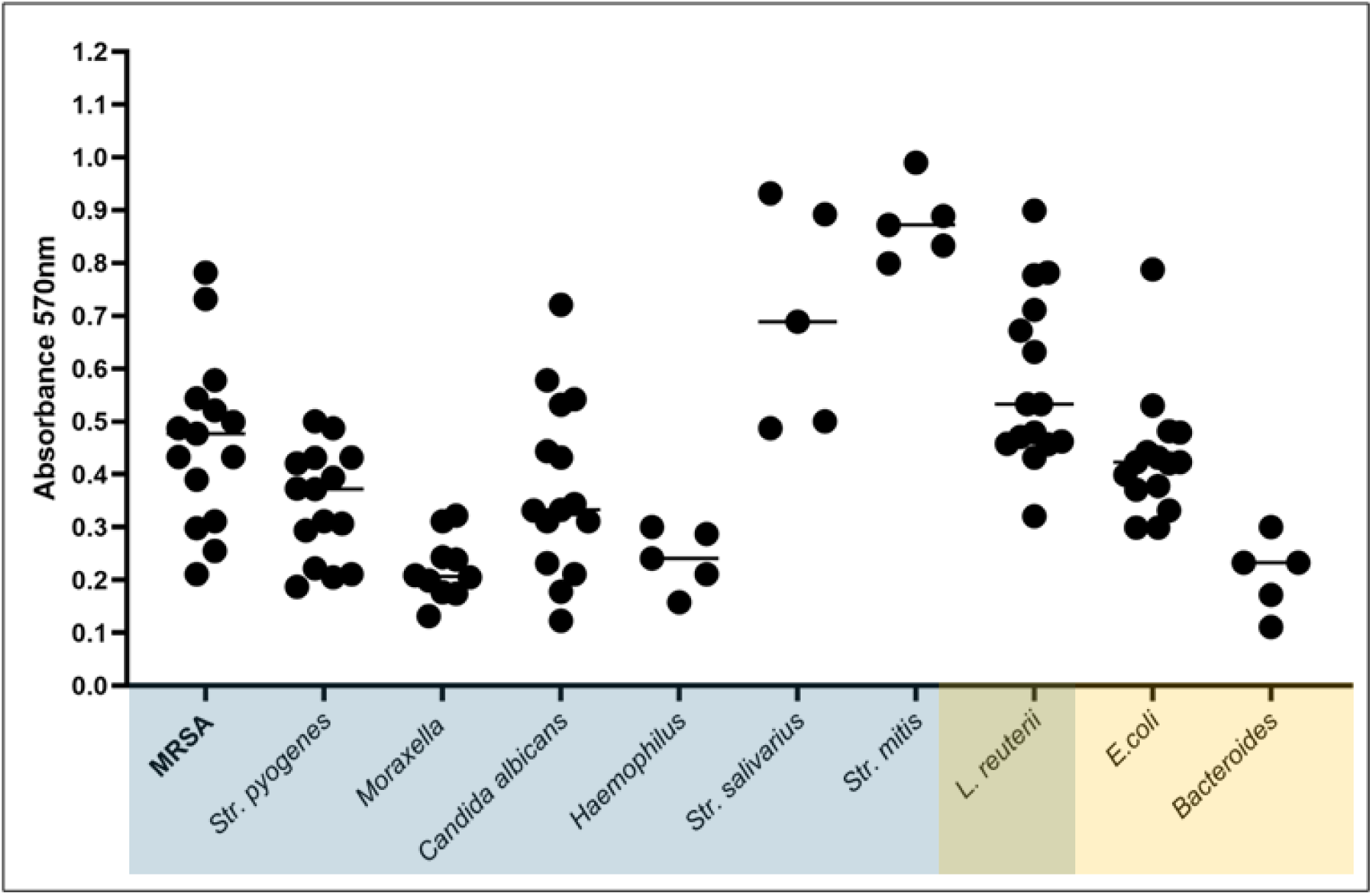
Determination of tested strains’ ability to form biofilm *in vitro* using crystal violet method. The blue box highlighted strains typical of oral cavity/throat, yellow box highlighted strains typical of intestine; both boxes overlap on *L. reuterii, i.e.* the probiotic strain found in both niches. The horizontal lines are median values. N=105.

**Figure 4.**
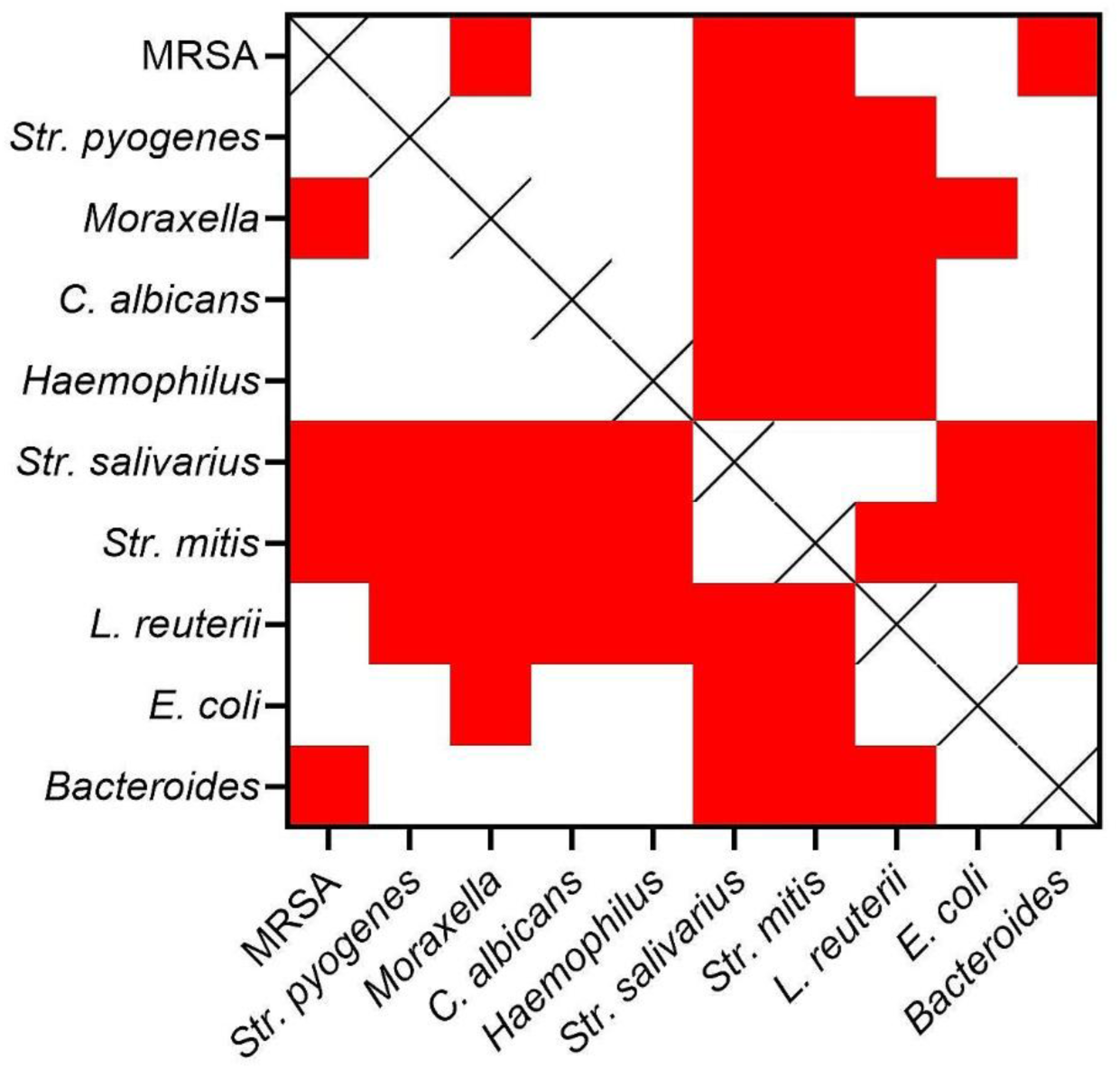
Comparison of statistical significances in strains’ ability to form biofilm. Statistical significance (p<0,001) marked as red squares; differences not statistically significant marked as white squares; X – comparison excluded.

Subsequently, for selected representative strains, the antimicrobial activity of pure octenidine dihydrochloride was compared with that of a clinically used formulation—octenidine-containing lozenges—at equivalent concentrations (**Table S1**). The values of MBC of octenidine in relevant contact times are presented in **Figure 5**. At the applied contact times and physiologically relevant concentrations, octenidine dihydrochloride exhibited strong biocidal activity against microorganism’s representative of the oral cavity and throat, with MBC values well below the achievable local concentration of 260 mg/L.

**Figure 5.**
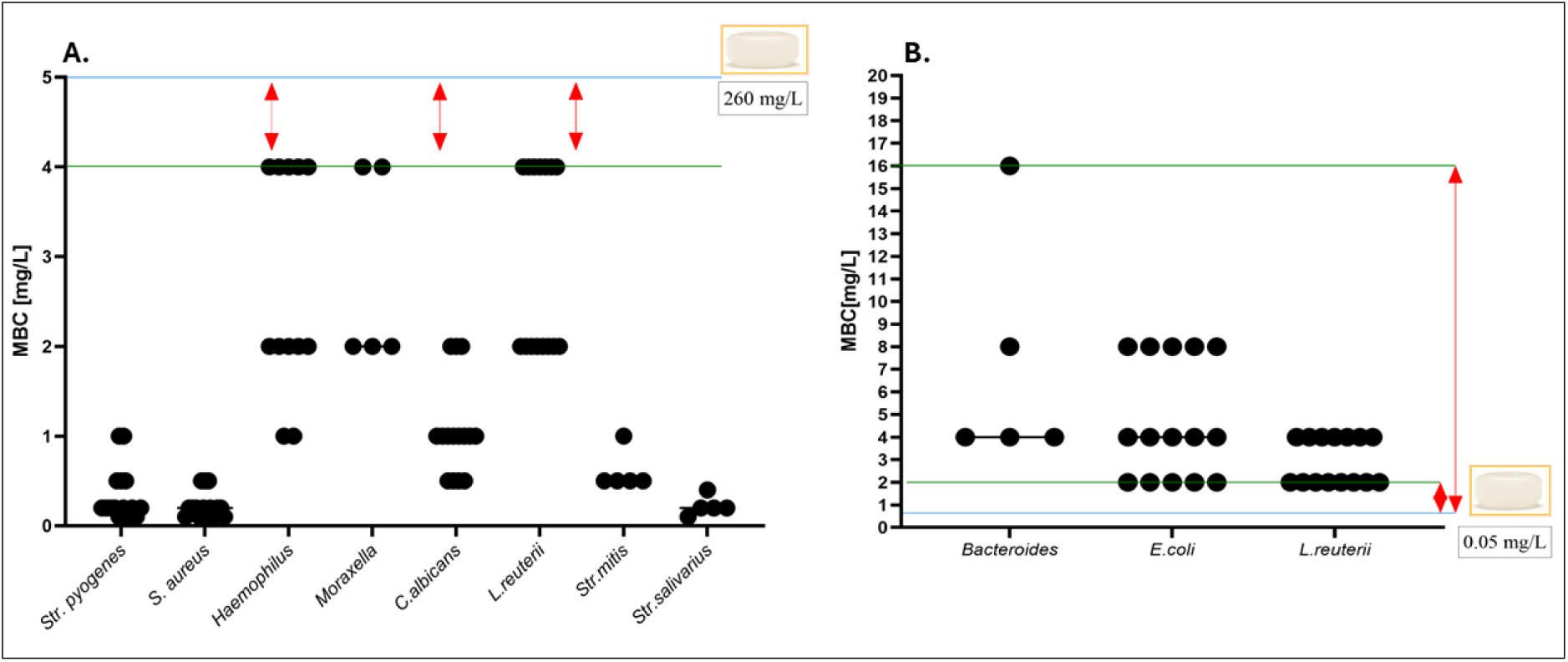
Minimal Biocidal Concentrations (MBCs) of octenidine dihydrochloride against the tested microorganisms. **(A)** MBC values determined for pathogens and commensals representative of the oral cavity and throat. **(B)** MBC values determined for strains representative of the intestinal microbiota. The blue line indicates the estimated maximum concentration of octenidine achievable in the respective anatomical region (260 mg/L for the oral cavity and 0.05 mg/L for the large intestine). The green line marks the highest (panel A) or lowest (panel B) MBC values observed within the tested group. In panel A, red arrows highlight the strains for which the highest MBC (4 mg/L) remains 65-fold lower than the locally achievable concentration of octenidine. In panel B, red arrows illustrate that the concentrations required to exert a biocidal effect toward intestinal commensals exceed the estimated in situ exposure by 4x-32-fold. N=105.

In contrast, no biocidal effect was observed against intestinal commensals at the estimated colonic concentration of 0.05 mg/L, as their MBC values exceeded this threshold by several orders of magnitude. In the next line of investigation, the antimicrobial activity of octenidine towards biofilms was performed (**Figure 6**).

**Figure 6.**
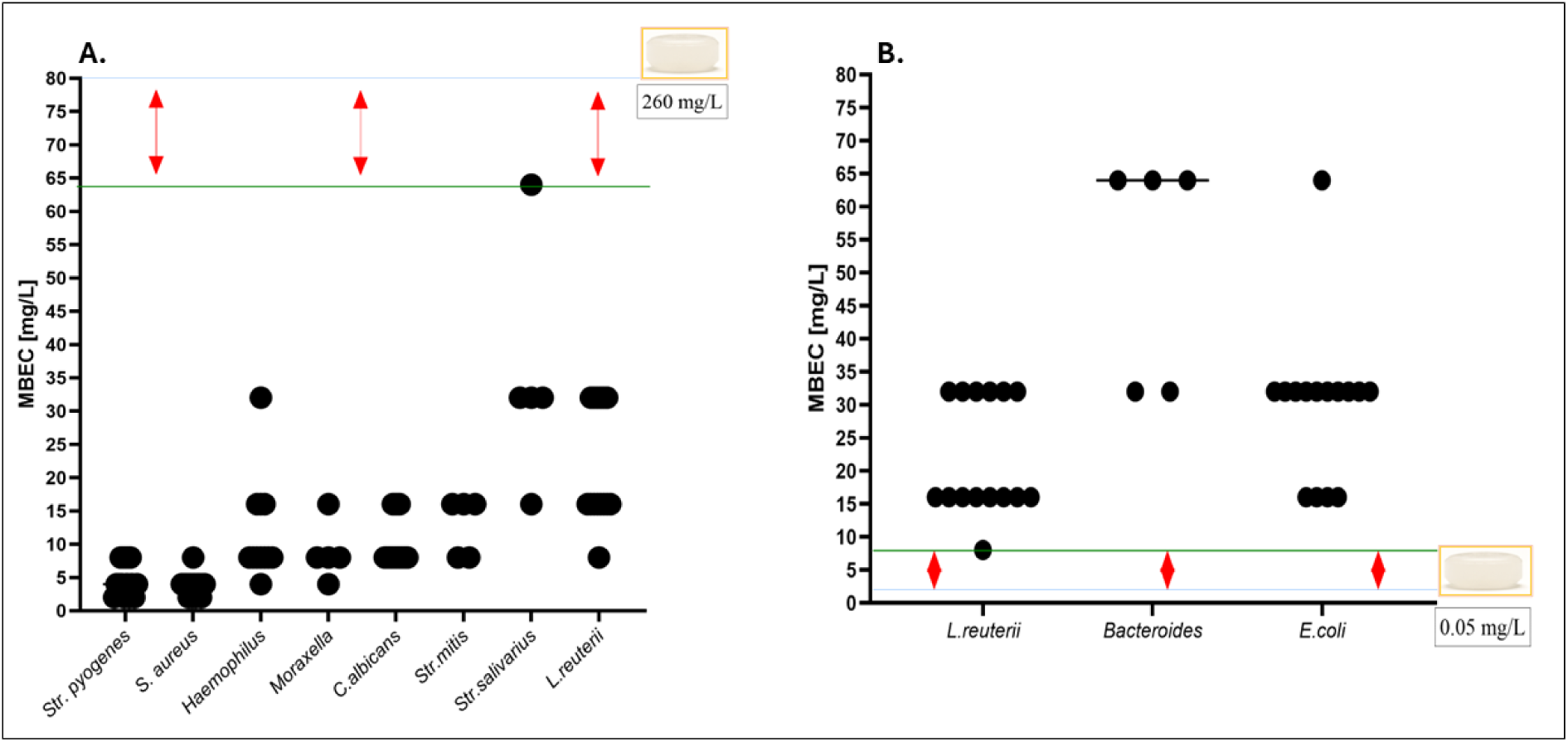
**(A)** MBEC values determined for pathogens and commensals representative of the oral cavity and throat. (**B**) MBEC values determined for strains representative of the intestinal microbiota. The blue line indicates the estimated maximum concentration of octenidine achievable in the respective anatomical region (260 mg/L for the oral cavity and 0.05 mg/L for the large intestine). The green line marks the highest (panel A) or lowest (panel B) MBEC values observed within the tested group. In panel A, red arrows indicate strains for which the lowest MBEC (8 mg/L) remains over 30-fold lower than the locally achievable concentration of octenidine. In panel B, red arrows show that the concentrations required to eradicate biofilms formed by intestinal commensals exceed the estimated *in situ* exposure by 8x–64x.

Under the applied exposure conditions, octenidine dihydrochloride was highly effective in eradicating biofilms formed by pathogens colonizing the oral cavity and throat, with MBEC values remaining below the local concentration of 260 mg/L. In contrast, biofilms formed by commensal strains isolated from the intestinal microbiota exhibited MBEC values significantly exceeding the estimated colonic concentration of 0.05 mg/L. N=105.

## 4. Discussion

Although antiseptics, such as octenidine dihydrochloride (**Figure 1**) are increasingly proposed as topical alternatives to antibiotics in the treatment of infections, due to its general non-specific antimicrobial mechanism of action (**Figure 2**, **Table 1**), little is known about their microbiological selectivity when administered orally. Lozenges containing octenidine dihydrochloride are specifically designed to act in the oral cavity and pharynx; however, their use inevitably results in swallowing of the dissolved antiseptic agent resulting in its potential action in the lower parts of the digestive tract including large intestine. This raises two critical and yet unresolved questions: (i) to what extent octenidine affects not only pathogenic but also commensal microorganisms at the desired site of application, and (ii) whether octenidine’s concentration reached in the large intestine may affect the intestinal microbiota. Despite octenidine’s long-standing use in oral rinses, no study to date has addressed this issue. This experiment was aimed at filling this gap.

In our previous work (14), we demonstrated that octenidine lozenges dissolve fully within an average of 23 minutes, releasing the active compound gradually into the saliva. This allowed us to construct a preliminary pharmacokinetic/pharmacodynamic model reflecting two distinct but physiologically relevant exposure scenarios: topical contact in the oral cavity and subsequent gastrointestinal passage after swallowing. The existing data on physicochemical properties of octenidine indicate that it is not systemically absorbed through the mouth mucosa or intestinal wall (23). Concentration of octenidine in saliva reaches approximately 260 mg/L in 23 minutes, the residual amount that could potentially reach the colon is estimated to be diluted to 0.05 mg/L over 24 hours. Although simple, this two-compartment model reflects to a certain extent the patient’s use of lozenges and provides a rationale for investigating microbial responses under these two defined exposure conditions.

All tested strains demonstrated the ability to form biofilm under *in vitro* conditions (**Figures 3**-**4**), which should be considered a normative trait rather than a virulence exception, particularly given the protected and nutrient-rich nature of mucosal niches (24). Nevertheless, we performed minimal biocidal concentration (MBC) and minimal biofilm eradication concentration (MBEC) testing to assess the antiseptic’s efficacy under both planktonic and sessile conditions (**Figure 5-6**). Octenidine, at the concentration typical to its presence in saliva (260 mg/L), exhibited potent activity against all tested oropharyngeal pathogens, including biofilm-forming *Str. pyogenes*, *S. aureus*, *C. albicans*, and *M. catarrhalis*. Contrarily, at the estimated colonic concentration (0.05 mg/L), no biocidal or biofilm-disrupting effect was observed against any tested representative of the intestinal microbiota, including obligatory anaerobes such as *Bacteroides spp*.

These findings suggest a high dependence of octenidine activity on both microbial strains’ susceptibility and concentration profiles. An important aspect our findings is the exact distinction between transient pathogens and resident commensals of the oral cavity and pharynx. While octenidine effectively eradicated pathogens, its impact on physiological microbiota—such as *Str. salivarius*, *Str. mitis*, and *L. reuterii*, although measurable, was of a size of a non-specific effect observed in the effect of routine oral hygiene practices, such as tooth brushing or antiseptic mouthwashes. Importantly, octenidine-based mouthwashes have been in wide use for over four decades, yet there are no reports on effecting dysbiosis. Oral hygiene always causes a transient reduction in microbial load, but this is a rapidly reversible phenomenon, supported by the regenerative potential of healthy microbiota and its synergistic interactions with the mucosa-associated lymphoid tissue (25). In this context, short-term exposure to antiseptics may be considered a protective function by tipping the ecological balance in favor of the host compared to pathogenic species, without causing lasting dysbiosis (26).

Another issue is whether swallowed octenidine, even in trace amounts, could negatively impact the gut microbiota. Our experimental findings do not support such possibility – none of the tested intestinal strains, including genera *Bacteroides* and *Escherichia*, was affected by octenidine at concentrations estimated to reach the colon. Importantly, our *in vitro* design deliberately excluded additionally protective physiological variables such as bile components, mucus, or interactions with food residues and stool matrix, which *in vivo* would further reduce antiseptic’s bioavailability and antimicrobial action (24). In that sense, our model was constructed to overestimate, rather than underestimate, potential intestinal impact, making the absence of any observed antimicrobial activity even more reassuring. This study is an exploratory investigation focused on establishing a physiologically grounded *in vitro* model to assess dual-compartment (oral cavity and large intestine) impact of orally administered octenidine. Of course, its simplified structure comes with certain limitations. Most importantly, the pharmacodynamic model used to simulate gastrointestinal exposure excluded aforementioned protective biological variables. In our understanding, incorporating more complex models in future studies - using artificial feces or dynamic gut simulators, would likely further attenuate the antiseptic’s residual activity in the intestinal environment, thereby reinforcing rather than weakening the conclusion that orally administered octenidine is microbiologically safe for the gut. Recognizing these limitations, we are currently planning a staged validation approach, beginning with a *Galleria mellonella* larval model, followed by *in vivo* studies in mice colonized with human-derived gut microbiota, to further explore host - microbe interactions under conditions that better mimic physiological reality.

In summary, our study provides a microbiological assessment of octenidine lozenges action, focusing on both their intended topical action in the oropharynx and potential unwanted effects in the rest of gastrointestinal tract. Using realistic exposure times and concentrations in our *in vitro* model, we demonstrated that octenidine effectively eradicates oropharyngeal pathogens, including biofilm-embedded forms, while exerting transient, “toothbrush-like” effects on commensals of the oral cavity and throat. Importantly, the estimated concentration reaching the large intestine proved safe to representative members of the physiological intestinal microbiota, even under conservative conditions. These findings prove that antiseptic lozenges containing octenidine offer targeted activity against pathogens in the upper digestive tract without compromising microbiota-associated homeostasis. As such, they may represent a rational, microbiologically safe alternative to systemic antibiotic therapy in selected clinical scenarios such as sore throat.

## 5. Conclusions

- octenidine dihydrochloride exhibits potent biocidal and antibiofilm activity against microorganisms residing in the oral cavity and pharynx.
- although the antiseptic also affects members of the physiological oral microbiota, this effect is comparable to the that during routine tooth brushing. As such, the rapid regenerative capacity of the normal oral microbiota and the transient nature of antiseptic exposure are expected to preserve long-term microbial homeostasis.
- octenidine did not affect any of the bacterial species of the large intestine. Intestinal microbiota displayed significantly higher MBC and MBEC values, far exceeding concentration of octenidine that could reach the colon after oral administration.
- octenidine, when delivered *via* oral tablets, provides targeted antimicrobial activity in the upper aerodigestive tract and is safe for intestinal microbiota.

