## Supplemental Table 1 for "Orally Administered Octenidine Is Effective Against Oral/Throat Pathogens and Safe for the Normal Microbiota of the Oral Cavity, Throat and Large Intestine"

| | Results compliance [Y/N] | |
|---|---|---|
| | OCT-lozenge | Pure OCT |
| *S. aureus 29213* | Y | Y |
| *P. aeruginosa 27853* | Y | Y |
| *C. albicans 10231* | Y | Y |

TabS1. Comparison of compliance of antimicrobial results of the same concentrations of octenidine from lozenges vs powder. Micro-plate method for MIC assessment; 24 hour contact/exposure time.
